# THA_1941 from *Thermosipho africanus*: A Thermostable β-1,3-Glucan Phosphorylase for Efficient β-1,3-Glucan Synthesis

**DOI:** 10.1101/2025.04.05.647330

**Authors:** Guotao Mao, Jin Yu, Junhan Lin, Ming Song, Zengping Su, Hui Xie, Hognsen Zhang, Hongge Chen, Andong Song

## Abstract

β-1,3-Glucan phosphorylases capable of utilizing glucose as a priming substrate are key biocatalysts for the synthesis of functional β-1,3-glucan. In this study, we identified THA_1941 from *Thermosipho africanus* (TaβGP) as a GH161 β-1,3-glucan phosphorylase exhibiting robust synthetic activity towards glucose, as confirmed by ^13^C nuclear magnetic resonance, liquid chromatography-mass spectrometry, and sequence and structural analyses. TaβGP displayed exceptional thermostability, retaining 93% of its activity at 60 °C for 180 h, and showed broad pH tolerance ranging from pH 5.0 to 10.0, surpassing the performance of previously reported homologs. In addition, TaβGP exhibited broad substrate flexibility, accepting both α- and β-linked disaccharides, and demonstrated strong resistance to metal ions and lignocellulose-derived inhibitors. In the presence of 150 mM glucose 1-phosphate as the donor substrate, TaβGP synthesized β-1,3-glucan with a tunable average degree of polymerization (10–32), depending on the concentration of glucose used as the primer. The combination of thermostability, inhibitor resistance, and substrate versatility makes TaβGP a promising biocatalyst for the economically viable and environmentally sustainable synthesis of β-1,3-glucan from non-food biomass sources.

## 1. Introduction

The production of value-added bioproducts from lignocellulosic biomass offers substantial economic and environmental advantages, supporting the development of a circular and sustainable economy(Chen et al., 2023). β-1,3-Glucan is a bioactive polysaccharide with diverse biological functions, including immunomodulation, antioxidation, anti-tumor, and anti-diabetic effects(Liu et al., 2021; Zeković et al., 2005). Consequently, β-1,3-glucan holds significant potential for applications in the pharmaceutical, cosmetic, food, and feed industries, with its global market projected to exceed USD 2.3 billion in the near future(Ahmad et al., 2024; Lehtovaara and Gu, 2011). Cereal and yeast currently serve as the primary sources of commercial β-1,3-glucan products. However, the low abundance of cell wall-associated β-1,3-glucan in plants and fungi necessitates complex and intensive extraction and purification processes, increasing production costs, posing environmental challenges, and resulting in structural heterogeneity, which ultimately limits its broader applications(Edo et al., 2024; Ma et al., 2021). Leveraging the low cost of agricultural biomass and by-products, combined with the strict specificity of enzymes, the bottom-up synthesis of β-1,3-glucan from agricultural wastes offers a sustainable alternative to enhance both the economic feasibility and structural homogeneity of β-1,3-glucan products.

*In vitro* synthetic biosystems, consisting of multiple enzymes catalyzing complex biotransformations, have merged as a promising biomanufacturing platform for β-1,3-glucan synthesis from cellobiose(Zhang, 2015). These biosystems offer several advantages, including high product yield, rapid reaction rates, precise control of reaction conditions, and scalability(Liu et al., 2023; Zhang, 2015). This platform has been successfully applied in the industrial production of inositol(You et al., 2017). In a designed *in vitro* synthetic biosystem of β-1,3-glucan from cellobiose, cellobiose phosphorylase (CBP) catalyzes the ATP-independent phosphorolysis of cellobiose into glucose-1-phosphate (G1P) and glucose in the presence of inorganic phosphate (P_i_)(Alexander, 1968). Subsequently, β-1,3-glucan phosphorylase synthesizes β-1,3-glucan using G1P as the donor substrate and glucose as the priming substrate(Kuhaudomlarp et al., 2019a). The strict regioselectivity of phosphorylases ensures the structural homogeneity of the synthesized β-1,3-glucan products, a critical feature for applications and structure-function studies(Kuhaudomlarp et al., 2019a). Industrial scalability depends heavily on the thermostability and catalytic efficiency of the phosphorylases employed. While the thermostable CBP from *Acetivibrio thermocellus* (CtCBP) has been successfully utilized in synthetic biosystems(Niu et al., 2025; You et al., 2013), suitable β-1,3-glucan phosphorylases remain limited.

Phosphorylases catalyzing β-1,3-glucan synthesis are classified into glycoside hydrolase (GH) families GH94, GH149, and GH161. Laminaribiose phosphorylase (LBP) from the GH94 family predominantly synthesizes laminaribiose using glucose as the priming substrate(Sun et al., 2024). β-1,3-Glucan phosphorylases in GH149 and GH161 exhibit distinct acceptor substrate specificities. GH149 phosphorylases synthesize β-1,3-glucan using glucose as the acceptor substrate(Kuhaudomlarp et al., 2018), whereas GH161 phosphorylases require the costly laminaribiose as the priming substrate and show no activity towards glucose(Kuhaudomlarp et al., 2019a). To date, only one thermostable β-1,3-glucan phosphorylase, discovered from *Anaerolinea thermophila* (AtβOGP), has been characterized(De Doncker et al., 2024). Previous studies reported THA_1941, a thermostable phosphorylase from *Thermosipho africanus*, exhibited synthetic activity toward glucose comparable to GH149 members EgP1 and Pro_7066, and catalyzed the phosphorolysis of cellobiose and cellodextrin(Kuhaudomlarp et al., 2018; Wu et al., 2017). However, conflicting reports indicated that THA_1941, as a GH161 β-1,3-glucan phosphorylase, did not phosphorolyze laminaribiose and exhibited limited synthetic activity with glucose as the acceptor substrate(Kuhaudomlarp et al., 2019a). These discrepancies necessitate further investigation into the potential of THA_1941 for industrial-scale β-1,3-glucan synthesis.

In this study, THA_1941 was identified as a thermostable and pH-stable β-1,3-glucan phosphorylase belonging to the GH161 family, demonstrating robust synthetic activity towards glucose. Furthermore, THA_1941 exhibited high tolerance to various metal ions present in cane molasses and lignocellulose-derived inhibitors, highlighting its strong potential for β-1,3-glucan synthesis from agricultural wastes.

## 2. Materials and methods

### 2.1. Reagents and recombinants plasmids

All chemicals were of reagent grade or higher and were purchased from Sinopharm (Shanghai, China), unless otherwise noted. G1P, glucose-6-phosphate (G6P), cellobiose, glucose, and sodium hexametaphosphate were obtained from Yuanye Biotechnology (China). Laminaribiose, laminaritriose, and laminaritetraose were sourced from Megazyme (Ireland). Silica gel 60 F_254_ aluminum sheets were supplied by Merck (Germany). Kanamycin, ampicillin, isopropyl β-D-1-thiogalactopyranoside (IPTG), and ammonium molybdate were purchased from Solarbio (China). G6P dehydrogenase (G6PDH) and glucose phosphate mutase (PGM) were obtained from Rhawn (China).

The pET21a-TaβGP plasmid overexpressing THA_1941 from *T. africanus* TCF52B (TaβGP, NCBI accession No.: ACJ76363.1) was preserved at Henan Agriculture University(Wu et al., 2017).

Site-directed mutagenesis was used to generate TaβGP mutants R345Q, R373Q, D625A, and E840Q. Primers used for mutagenesis were listed in Table S1. Polymerase chain reactions were conducted with pET21a-TaβGP as the template. The resulting PCR products were treated with the restriction enzyme *Dpn* I at 37 °C for 3 h and transformed into *Escherichia coli* DH5α cells. Mutations were confirmed via DNA sequencing.

### 2.2. Expression and purification of recombinant enzymes

*E. coli* BL21(DE3) cells harboring the recombinant plasmids were cultured in LB medium supplemented with 100 μg/mL ampicillin or 50 μg/mL kanamycin at 37 °C. Protein expression was induced with 0.5 mM IPTG for 16 h at 16°C when the optical density at 600 nm reached 0.8. All purification procedures were performed at 4°C. Harvested cells were lysed using a JN-Mini homogenizer (China), and recombinant enzymes in the supernatant were purified via Ni-NTA affinity chromatography(Mao et al., 2021). The purity of the purified enzymes was analyzed by SDS-PAGE, and protein concentrations were measured using a bicinchoninic acid (BCA) protein assay kit (NCM Biotech, China). Purified enzymes in 50 mM Tris-HCl (pH 7.5) were stored at −80 °C for subsequent experiments.

### 2.3. Activity assays of recombinant enzymes

The activities of TaβGP in both synthetic and phosphorolytic reactions were evaluated. In the synthetic reaction, one unit (U) of TaβGP activity was defined as the amount of enzyme required to release 1 μmol of P_i_ per minute. Appropriate diluted TaβGP was added to the reaction mixture containing 50 mM sodium citrate buffer (pH6.0), 40 mM G1P, 40 mM glucose, and 5 mM MgCl_2_. The reaction was incubated at 50 °C for 10 min, then terminated by boiling and cooled to room temperature. Ammonium molybdate and ascorbic acid were subsequently added for color development at 30 °C for 15 minutes. The absorbance at 850 nm was measured to determine P_i_ concentration(De Groeve et al., 2010). The phosphorolytic activity of TaβGP was determined using a previously reported method(Nihira et al., 2012). One unit (U) of TaβGP activity was defined as the amount of enzyme required to produce 1 μmol of G1P per minute. The reaction mixture contained 50 mM sodium citrate buffer (pH 6.0), 1 mM laminaribiose, 100 mM NaH_2_PO_4_, and appropriately diluted TaβGP. After incubation at 50 °C for 20 minutes, the reaction was terminated by boiling. G1P content was determined by adding 50 mM Tris-HCl buffer (pH 7.5), 3 mM NADP^+^, 3 U/mL G6PDH, and 3 U/mL PGM. After incubation at 30 °C for 20 min, absorbance was measured at 340 nm to calculate NADPH generated from G1P.

### 2.4. Effects of temperature and pH on TaβGP activity

The effects of temperature and pH on the activity and stability of TaβGP were evaluated using its activity in the synthetic reaction, given the enzyme’s critical role in β-1,3-glucan synthesis.

The effect of temperature on TaβGP activity was determined in 50 mM sodium citrate buffer (pH6.0) at temperature ranging of 40 °C to 90 °C. For thermostability evaluation, TaβGP were incubated at different temperatures (50–80 °C) for various time intervals, and residual activity was measured.

The optimal pH for TaβGP was determined using 50 mM sodium citrate buffer (pH 4.0–6.0) and 50 mM Tris-HCl buffer (pH 7.0–10.0) at 60 °C. To assess pH stability, TaβGP were incubated in buffers spanning pH 5.0–10.0 on ice for 8 h, and residual activity was subsequently measured.

### 2.5. The kinetics of TaβGP

The kinetics of the synthetic reaction were assessed using 40 mM G1P, and varying concentrations of glucose (1–50 mM), or laminaribiose, laminaritriose, and laminaritetraose (0.2–5 mM) at 60 °C in 50 mM sodium citrate buffer (pH 6.0). The kinetic parameters, including *k*_cat_ and *K*_m_, were calculated by fitting data to the Michaelis-Menten equation using GraphPad Prism 8.

### 2.6. Substrate specificity of TaβGP

The synthetic activity of TaβGP on various acceptor substrates was determined in reactions containing 50 mM G1P, appropriately diluted TaβGP, and 1 mM various glucose-glucose (Glc-Glc) disaccharides or monosaccharides at 50 °C. For phosphorolytic reaction, the cleavage of the various Glc-Glc disaccharides was carried out in the presence of 10 mM NaH_2_PO_4_. The tested monosaccharides included glucose, glucosamine (GlcN), N-acetyl-glucosamine (GlcNac), 2-deoxy-glucose (DGlc), xylose, mannose, and arabinose. The tested Glc-Glc disaccharides were trehalose (α-1,1), kojibiose (α-1,2), nigerose (α-1,3), maltose (α-1,4), isomaltose (α-1,6), sophorose (β-1,2), laminaribiose (β-1,3), cellobiose (β-1,4), and gentiobiose (β-1,6). Cellotriose and laminaritriose were also evaluated as substrates.

Reactions were terminated by boiling and analyzed using thin-layer chromatography (TLC) on Silica Gel 60 F_254_ plates developed in n-butanol:acetic acid:water (2:1:1). Bands were visualized using sulfuric acid:methanol (1:4) staining and heating at 120 °C..

For the phosphorolysis of cellobiose, cellotriose, laminaribiose, and laminaritriose, the G1P product was further analyzed by liquid chromatography-mass spectrometry (LC-MS) using an Agilent 1290II-6460 system in negative ion mode. Chromatographic separation was performed with a C18 analytical column (BEH-C18, 2.1 × 100 mm, 1.7 μm, Waters) under gradient elution. The mobile phases were 0.1% formic acid in water (eluent A) and 0.1% formic acid in acetonitrile (eluent B), with a flow rate of 0.25 mL/min. The gradient was as follows: 0–4–8–12–14–20 min, 1–1%– 10%–80%–80%–1% B.

### 2.7. Effects of metal ions and organic solvents on the activity of TaβGP

The influence of metal ions and organic solvents on the synthetic activity of TaβGP was determined at 50 °C under the conditions of pH 6.0, 1 mM glucose, 40 mM G1P and different tested additives. Ethylenediaminetetraacetic acid (EDTA, 50 mM), metal ions (50 mM) including Na^+^, K^+^, Ca^2+^, and Mg^2+^ and organic solvents including furfural, 5-hydroxymethylfurfural (5-HMF), vanillin, and 4-hydroxybenzaldehyde (4-HBA) were assessed. Metal ions mixture (Na^+^, K^+^, Ca^2+^, and Mg^2+^, 50 mM each), and organic solvents mixture (furfural, 5-HMF, vanillin, and 4-HBA, 20 or 50 mg/L each) were also tested. The control samples were assessed in parallel without the addition of metal ions or organic solvents.

### 2.8. Structural characterization of β-1,3-glucan

The β-1,3-glucan synthesis from glucose and G1P was conducted at 50 °C for 6 h under the conditions of 150 mM G1P, 1–50 mM glucose, and 2 U/ml TaβGP. The reaction was terminated by boiling and subsequently cooled to room temperature. The resulting insoluble β-1,3-glucan products were precipitated by centrifugation and subsequently analyzed using ^13^C nuclear magnetic resonance (NMR), X-ray diffraction (XRD), Fourier transform infrared spectroscopy (FTIR), and scanning electron microscopy (SEM). For further characterization, β-1,3-glucan was precipitated using 90%(v/v) ethanol at 4°C, washed, and dried by vacuum freeze-drying. The degree of polymerization (DP) was determined using the matrix assisted laser desorption/ionization–time-of-flight mass spectrometry mass spectrometry (MALDI-TOF MS).

The synthesized β-1,3-glucan was dissolved in dimethyl sulfoxide (DMSO)-d_6_ and analyzed using ^13^C NMR spectroscopy on a Bruker AVANCE III 400M NMR spectrometer. Chemical shifts (δ) were recorded in ppm with solvent peaks used as internal references. The synthesized β-1, 3-glucan was subjected to the Bruker D8 ADVANCE X-ray diffractometer. XRD analysis was conducted using a Bruker D8 ADVANCE diffractometer, with XRD patterns recorded over a 2*θ* range of 2° to 40° in 0.02° increments, utilizing Cu Kα radiation (λ = 1.5418 Å) at 40 kV and 40 mA. FTIR spectra were obtained using a Thermo-Nicolet 6700 FTIR spectrometer over a spectral range of 400–4000 cm^−1^. Glucan 300, a commercial β-1,3-glucan product extracted from *Saccharomyces cerevisiae* (Transfer Point, Columbia, USA) with significant biological activities was selected as a reference(Vetvicka and Vetvickova, 2010). SEM imaging was performed using a Thermo Fisher Apreo 2 SEM equipped with a solid-state ultrafast UV laser (Nd:YAG, 355 nm). Dried samples were dispersed onto copper grids coated with a carbon support film and sputter-coated with gold before imaging at an accelerating voltage of 20 kV.

The molecular weight distribution of the synthesized β-1,3-glucan was determined using a MALDI-709 TOF MS instrument following the method described by (Petrović et al., 2015). The number-average molecular weight 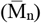, weight-average molecular weight 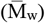, polydispersity index (PDI), and average degree of polymerization (DP_n_) were calculated using the corresponding equations.

### 2.10. Sequence and structure analyses of TaβGP

The protein sequences of selected members of the GH94, GH149, and GH161 families were aligned by MUSCLE in MEGA11, and then the phylogenetic tree was constructed by using the bootstrap value with 1000 replicates using the neighbor-joining method(Tamura et al., 2021). The structure of TaβGP binding with G1P was predicted using Protenix server(Team et al., 2025). Structural analyses of the TaβGP and its complexes with G1P were performed using PyMol(DeLano, 2002).

## 3 Results and discussions

### 3.1 Identification of THA_1941 as a β-1,3-glucan phosphorylase

#### 3.1.1 Sequence and structure analyses of THA_1941

Previous studies have indicated that THA_1941 from *T. africanus* may function as a thermostable phosphorylase, with significant potential for the production of value-added β-1,3-linked or β-1,4-linked glucan(Kuhaudomlarp et al., 2019a; Wu et al., 2017). To confirm the enzymatic identity of THA_1941, its sequence and structural characteristics were analyzed (Fig. 1).

**Fig. 1.**
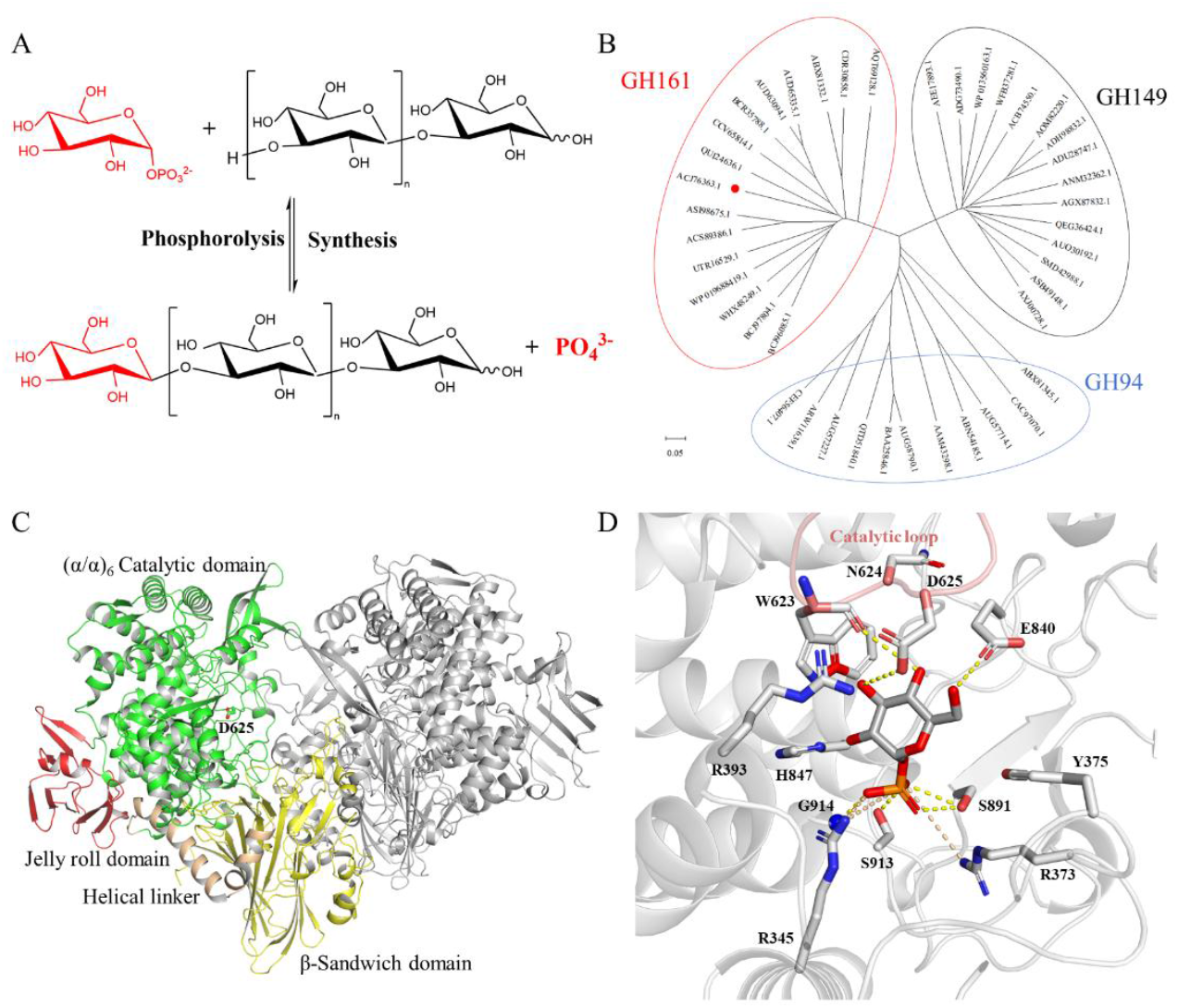
Sequence and structure analyses of THA_1941. (A) Schematic illustration of the reversible phosphorolytic reaction catalyzed by β-1,3-glucan phosphorylase. n≥0. (B) Phylogenetic tree of THA_1941. The position of THA_1941was highlighted with a red dot. (C) Overall structure of THA_1941. The catalytic residue D625 was showed as sticks. (D) Interaction of THA_1941 with G1P. Hydrogen bonds and ions interactions were labeled with yellow and wheat dashed lines, respectively.

Phylogenetic analysis revealed that THA_1941 belongs to the GH161 family, distinct from the GH149 and GH94 families (Fig. 1B). Sequence alignment demonstrated that THA_1941 shared 20.5% and 44.9% sequence identity with the characterized GH149 AtβOGP and GH161 PapP, respectively, and retained conserved catalytic and substrate-binding residues (Fig. S1). Structurally, THA_1941 exhibited a similar fold to the structure-solved GH149 Pro_7066, comprising an N-terminal β-sandwich domain, a helical linker, an (α/α)_6_ catalytic domain, and a C-terminal jelly roll domain(Kuhaudomlarp et al., 2019b) (Fig. 1C). The donor substrate G1P was positioned within the active site of THA_1941, showing interactions with conserved residues (Fig. 1D). Specifically, residues W623, N624, D625 within the catalytic loop, and E840 stabilized the glucose moiety of G1P through hydrogen bonding. Additionally, residues R345, R373, S891, and S913 interacted with the phosphate group of G1P via hydrogen bonds and ionic interactions. The binding mode of G1P in THA_1941 closely resembled those observed in GH149 Pro_7066 and GH94 LBP from *Paenibacillus* sp.(Kuhaudomlarp et al., 2019b; Kuhaudomlarp et al., 2019c). In summary, sequence and structure analyses confirmed that THA_1941 is a member of the GH161 family and suggested that it has potential activity toward β-1,3-glycoside substrates.

#### 3.1.2 Reversible cleavage of β-1,3-linked glycosides catalyzed by THA_1941

To validate the findings from the sequence and structure analyses, the enzymatic function of THA_1941 was investigated with the purified recombinant protein (Fig. S2A). Mutations in the conserved residues R345, R373, D625, and E840 disrupted interactions with G1P, leading to the inactivation of THA_1941 variants (Fig. S2B), which corroborated the results of the structural analysis.

β-1,3-Glucan phosphorylase catalyzes the revisable phosphorolysis of β-1,3-glycosides (Fig. 1A). In the phosphorolysis direction, substrates including β-1,4-linked cellobiose and cellotriose, as well as β-1,3-linked laminaribiose and laminaritriose, were tested in the presence of P_i_. TLC analysis revealed that THA_1941 phosphorolyzed laminaribiose and laminaritriose, but not cellobiose or cellotriose (Fig. 2A). These findings were confirmed by LC-MS, which demonstrated THA_1941’s specific activity on β-1,3-linked glycosides (Figs. 2C, 2D, and S3). Additionally, α-linked disaccharides and other β-linked (β-1,2 and β-1,6) disaccharides were not cleaved by THA_1941 (Fig. S4).

**Fig. 2.**
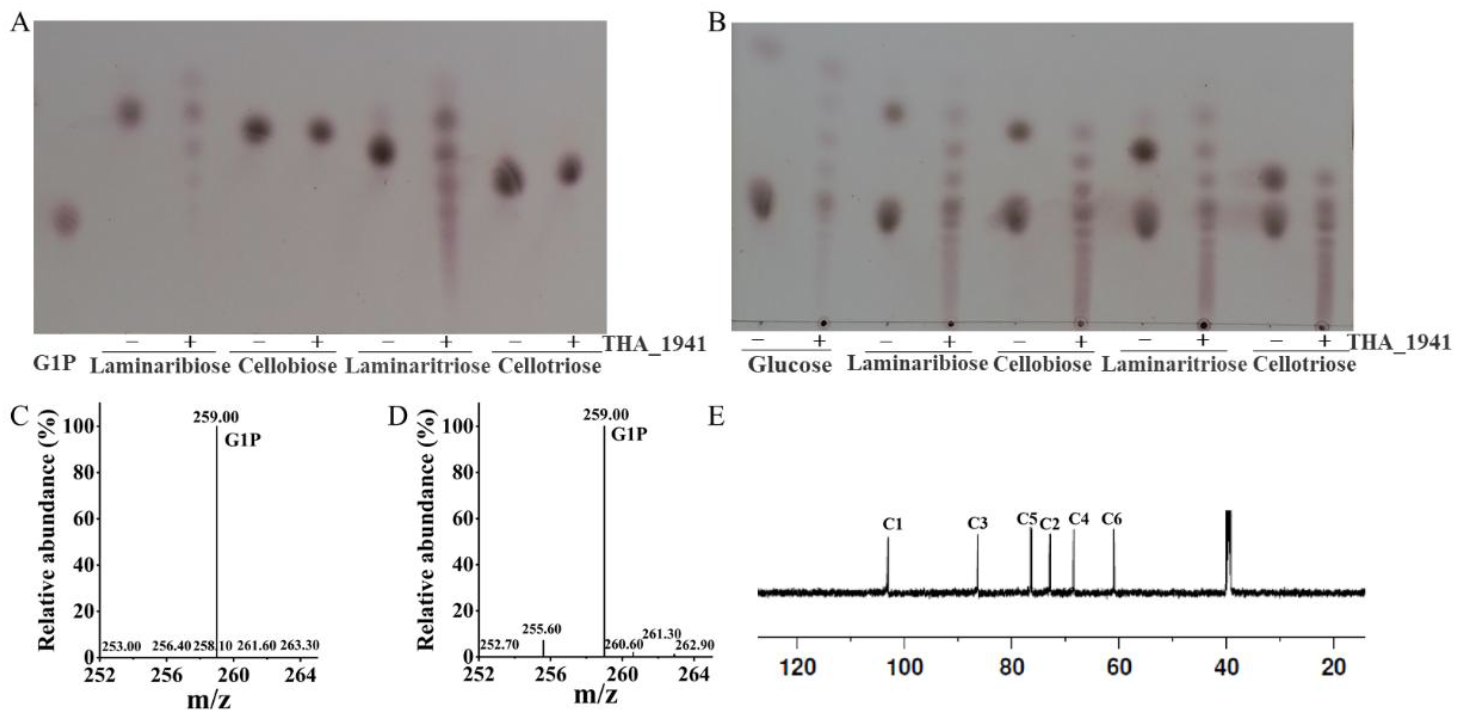
Identification of THA_1941 as a β-1,3-glucan phosphorylase. TLC analysis of the phosphorolytic (A) and synthetic (B) reaction catalyzed by THA_1941. Detection of G1P produced in the phosphorolysis of laminabiose (C) and laminaritriose (D) catalyzed by THA_1941. (E) ^13^C NMR analysis of the synthesized products catalyzed by THA_1941 using glucose and G1P.

In the synthetic direction, THA_1941 demonstrated the ability to utilize glucose, cellobiose, cellotriose, laminaribiose, and laminaritriose as acceptor substrates to synthesize glucans, with glucose-1-phosphate (G1P) serving as the donor substrate (Fig. 2B). The glucan synthesized from glucose and G1P was further characterized by ^13^C NMR spectroscopy, with the observed chemical shifts aligning with the reference spectra for linear β-1,3-glucan(Zheng et al., 2016) (Fig. 2E and Table S2). Notably, THA_1941 exhibited effective phosphorolytic activity toward β-1,3-glucans of any degree of polymerization (Fig. 1A, n ≥ 0), a feature distinct from the previously characterized GH161 PapP, which acted on β-1,3-glucans with n ≥ 1(Kuhaudomlarp et al., 2019a). This unique reversible activity allows THA_1941 to synthesize β-1,3-glucan directly from cost-effective glucose as the priming substrate, eliminating the need for expensive laminariobiose (Fig. 1).

Functional characterization conclusively identified THA_1941 from *T. africanus* (hereafter referred to as TaβGP) as a β-1,3-glucan phosphorylase belonging to the GH161 family, showing great promise for the economical synthesis of β-1,3-glucan using glucose as the priming substrate.

### 3.2 Characterization of TaβGP

To assess the industrial potential of TaβGP for β-1,3-glucan synthesis, the enzyme’s activity under varying temperature and pH conditions was evaluated.

The optimal temperature for TaβGP activity was determined to be 60 °C, with the enzyme retaining more than 95% of its maximum activity within the 50–60 °C range (Fig. 3A). TaβGP also displayed a broad pH optimum, with maximal activity between pH 6.0 and 7.0, demonstrating robustness across varying pH conditions (Fig. 3B). Furthermore, the enzyme retained 95% of its activity after incubation in buffers with pH values ranging from 5.0 to 10.0, confirming its pH stability (Fig. 3D).

**Fig. 3.**
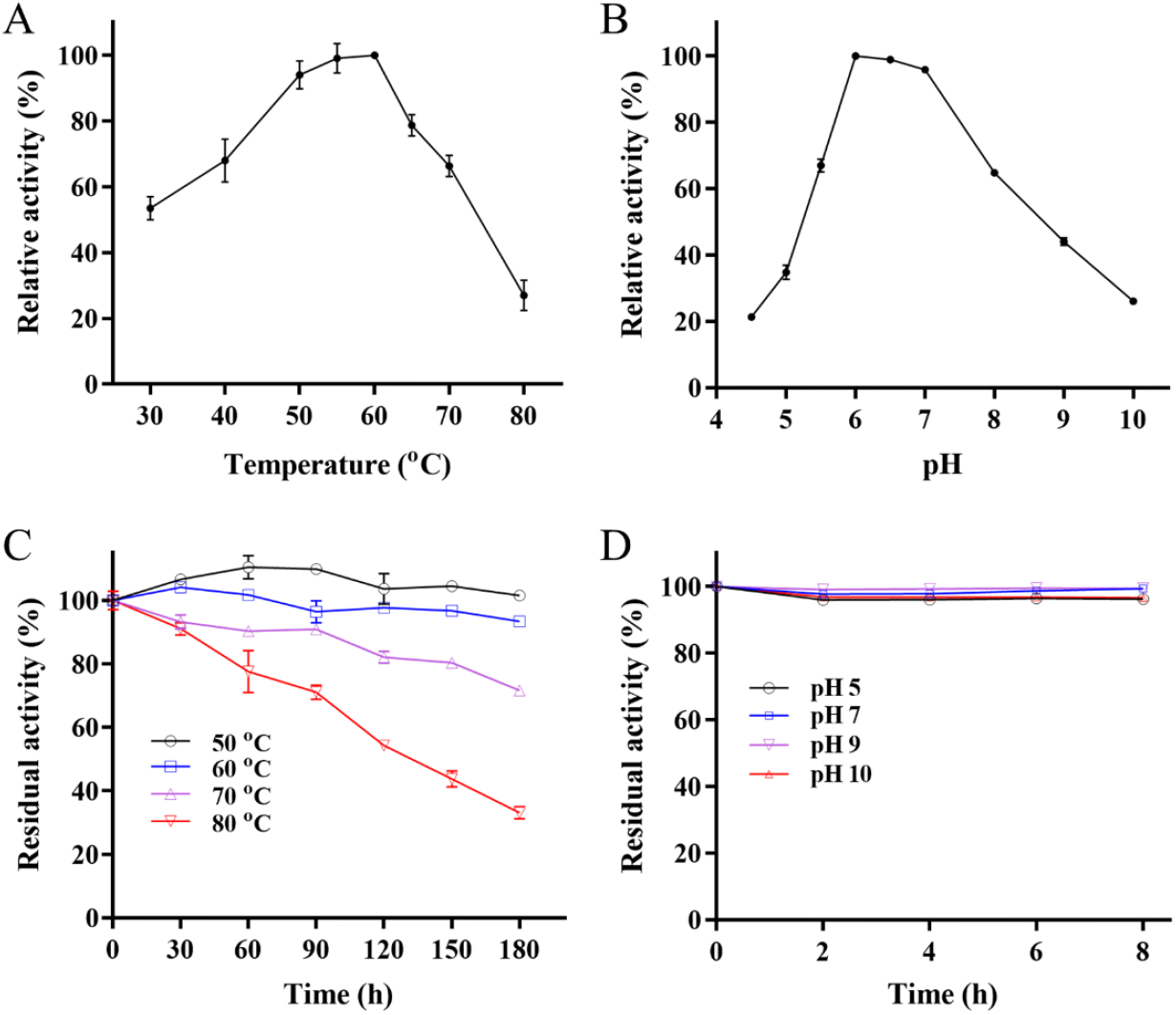
Effects of temperature and pH on the activity and stability of TaβGP. The optimal temperature (A) and pH (B) of TaβGP. The thermal (C) and pH (D) stability of TaβGP.

Thermal stability, a critical parameter for assessing industrial applicability, was a standout feature of TaβGP. After 180 h of incubation, TaβGP retained 100% of its original activity at 50 °C and 93% at 60 °C (Fig. 3C). Remarkably, TaβGP exhibited a half-life exceeding 120 h at 80°C. In contrast, other characterized β-1,3-glucan phosphorylases, such as EgP1 and Pro_7066, completely lost activity after incubation at 60°C for just 0.5 h(Kuhaudomlarp et al., 2018). Similarly, the characterized thermostable AtβOGP lost 30% of its activity after incubation at 60°C for 5 h(De Doncker et al., 2024). To our knowledge, TaβGP surpasses all previously reported β-1,3-glucan phosphorylases in terms of thermostability.

Table 1 illustrated TaβGP’s catalytic efficiency (*k*_cat_/*K*_m_), which was highest for laminaribiose and declined with increasing substrate chain length up to laminaritetraose. The turnover number (*k*_cat_) of TaβGP for laminaribiose was 11-, 339-, and 237-fold higher than those of PapP, EgP1, and Pro_7066, respectively(Kuhaudomlarp et al., 2018; Kuhaudomlarp et al., 2019a). A key factor in the economic feasibility of β-1,3-glucan production is the ability to synthesize it using glucose as the priming substrate. While the catalytic efficiency of TaβGP, EgP1, and Pro_7066 for glucose were comparable, TaβGP demonstrated a turnover number for glucose of 56.6 s^−1^, which was 51- and 34-fold higher than those of EgP1 and Pro_7066, respectively(Kuhaudomlarp et al., 2018). In contrast, GH161 family PapP showed no detectable activity towards glucose(Kuhaudomlarp et al., 2019a). The kinetic data suggested that while laminaribiose formation from G1P and glucose is the rate-limiting step, TaβGP efficiently catalyzes the subsequent synthesis of β-1,3-glucan once this intermediate is formed.

**Table 1.** Kinetic parameters of TaβGP in the synthetic direction.

| acceptor | $K_m$ (mM) | $k_{cat}$ (s<sup>-1</sup>) | $k_{cat}/K_m$ (s<sup>-1</sup>mM<sup>-1</sup>) |
| --- | --- | --- | --- |
| Glucose | 19.2±1.2 | 56.6±0.5 | 2.9 |
| Laminaribiose | 2.1±0.2 | 365.8±0.2 | 173.8 |
| Laminaritriose | 1.5±0.6 | 182.0±0.2 | 118.0 |
| Laminaritetraose | 1.3±0.1 | 164.1±0.3 | 131.0 |

In summary, TaβGP is a highly thermostable and pH-stable β-1,3-glucan phosphorylase. Its ability to utilize glucose as a priming substrate and its unmatched thermostability make it a promising candidate for the cost-effective production of β-1,3-glucan at an industrial scale.

### 3.3 Acceptor specificity of TaβGP

To broaden the range of substrates and enhance the diversity of synthesized glucans, the acceptor specificity of TaβGP was investigated using TLC analysis.

TaβGP was capable of synthesizing glucans using both α-linked and β-linked Glc-Glc disaccharides as acceptors, with G1P serving as the donor substrate (Fig. 4A). This contrasts with PapP in GH161 family and those in GH149 family, which showed no detectable activity toward α-linked disaccharides(Kuhaudomlarp et al., 2019a). The mechanism underlying TaβGP’s unique ability to utilize α-linked disaccharides remains to be elucidated.

**Fig. 4.**
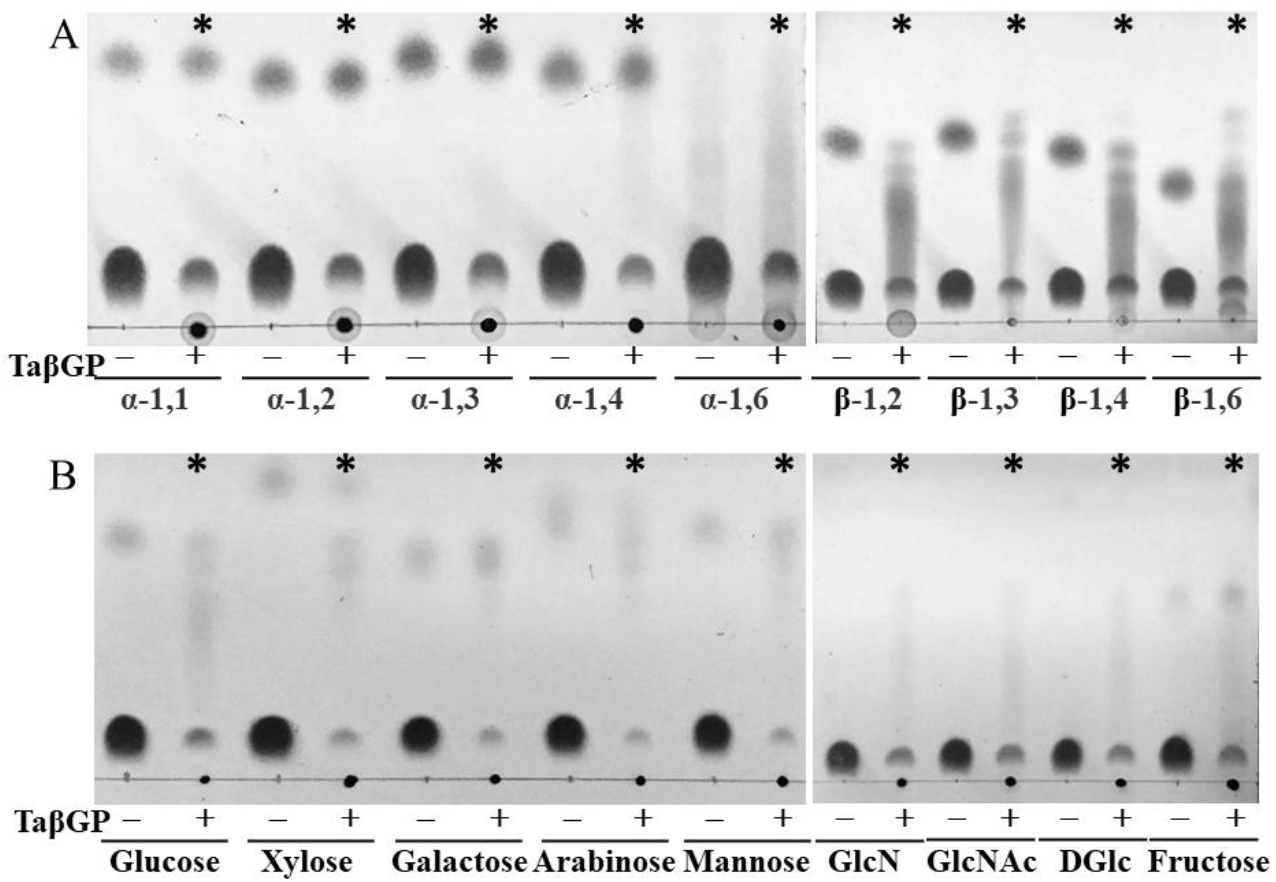
Substrate specificity of TaβGP. TLC analysis of the synthetic reaction catalyzing TaβGP with Glc-Glc disaccharides (A) and monosaccharides (B) as the priming substrate. The linkages of the disaccharides were indicated. GlcN, GlcNAc and DGlc represented D-Glucosamine, N-Acetyl-D-glucosamine, 2-Deoxy-D-glucose, respectively. The effective substrates of TaβGP detected by TLC were marked with asterisks.

In addition to disaccharides, TaβGP demonstrated activity with a variety of monosaccharides, including fructose, xylose, and arabinose, as acceptor substrates (Fig. 4B). Furthermore, TaβGP effectively utilized modified glucose derivatives such as glucosamine, N-acetyl-D-glucosamine, and 2-deoxy-D-glucose (Fig. 4B).

The ability of TaβGP to utilize a wide range of acceptor substrates, including both α-linked disaccharides and diverse monosaccharides, not only distinguishes it from other glycoside phosphorylases but also underscores its potential for creating structurally diverse β-1,3-glucans with enhanced functionalities.

### 3.4 Effects of different metal ions and organic solvents on the synthetic activity of TaβGP

The synthesis of value-added glycosides from agricultural waste, such as cane molasses and lignocellulose, presents a cost-effective and environmentally friendly alternative(Du et al., 2022; Xu et al., 2024; You et al., 2013). However, metal ions and organic solvents in these raw materials can inhibit enzyme activity. Here, the effects of metal ions and lignocellulose-derived inhibitors on the catalytic performance of TaβGP were investigated.

Cane molasses typically contain metal ions such as Na^+^, K^+^, Ca^2+^, and Mg^2+^(Xu et al., 2024). As showed in Fig. 5A, TaβGP’s synthetic activity was stimulated by Na^+^ and K^+^ ions, with activity increasing to 110% and 107%, respectively. Additionally, TaβGP showed no significant loss of activity in the presence of 50 mM Ca^2+^, and Mg^2+^. Furthermore, even when Na^+^, K^+^, Ca^2+^, and Mg^2+^ (each at 50 mM) were added simultaneously, TaβGP retained 110% of its original activity (Fig. 5A), indicating the high tolerance of TaβGP to the main metal ions typically present in the cane molasses. These results demonstrated that TaβGP, when coupled with sucrose phosphorylase, could efficiently catalyze β-1,3-glucan synthesis using sucrose derived from cane molasses, a by-product of the sugar industry.

**Fig. 5.**
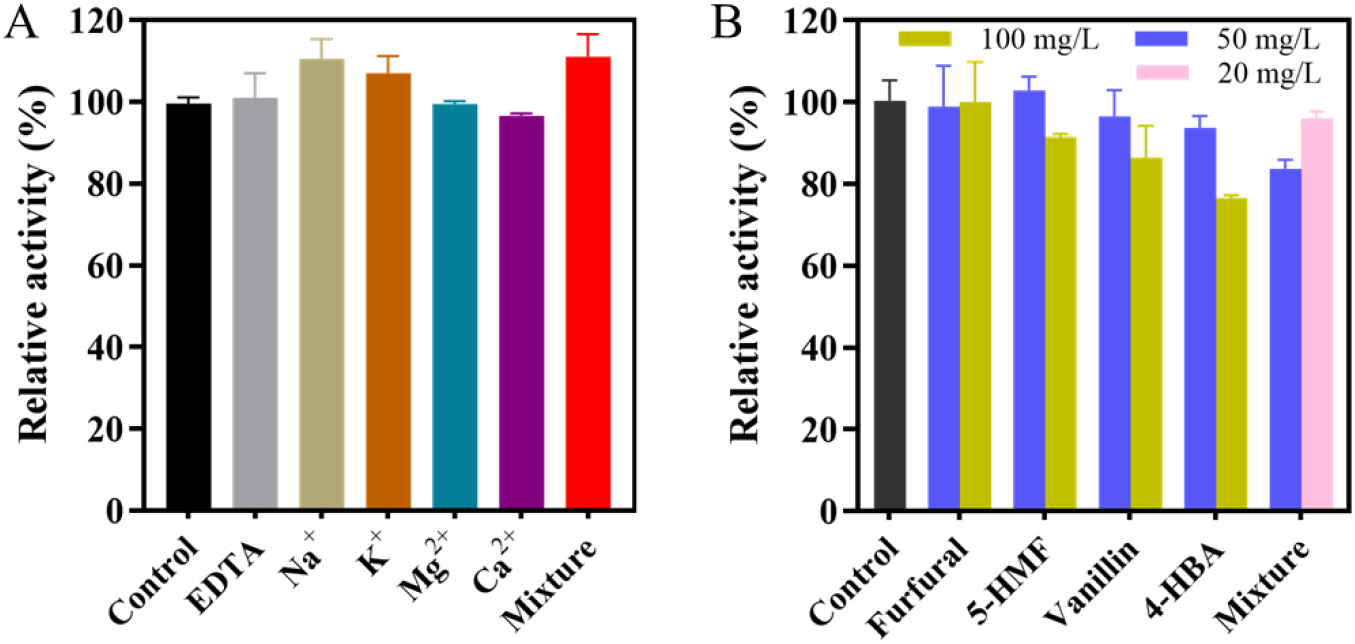
Effects of metal ions in cane molasses (A) and lignocellulose-derived inhibitors (B) on the synthetic activity of TaβGP. The mixture in the panel A and B represented the combined presence of Na^+^, K^+^, Ca^2+^, and Mg^2+^, and the combined presence of furfural, 5-HMF, vanillin, and 4-HBA, respectively.

Pretreatment of lignocellulose generates various inhibitors, including furfural, 5-HMF, vanillin, and 4-HBA, which are known to inhibit cellulases and other microbial enzymes(Guo et al., 2022; Jönsson and Martín, 2016; Wang et al., 2020). TaβGP demonstrated notable resistance to these inhibitors (Fig. 5B). At 50 mg/L, furfural, 5-HMF, and vanillin caused no observable inhibition. For 4-HBA, TaβGP retained 93% of its activity at 50 mg/L and 77% at 100 mg/L. Given that 4-HBA typically occurs at concentrations of less than 5 mg/L in lignocellulose hydrolysates(Zhang et al., 2013), TaβGP’s resilience to this inhibitor is highly promising. When exposed to combinations of furfural, 5-HMF, vanillin, and 4-HBA at 20 mg/L or 50 mg/L each, TaβGP activity decreased by only 4% and 10%, respectively (Fig. 5B). This demonstrated the TaβGP’s robustness and high tolerance to multiple inhibitors simultaneously, further supporting that its suitability for β-1,3-glucan synthesis from agricultural waste hydrolysates without detoxification steps.

TaβGP’s high tolerance to metal ions and lignocellulose-derived inhibitors solidifies its potential as a robust enzyme for β-1,3-glucan synthesis from agricultural waste. Its ability to operate effectively under challenging conditions transforms these wastes into valuable resources, contributing to sustainable and economically viable industrial processes.

### 3.5 Characterization of the synthesized β-1,3-glucan by TaβGP using glucose and G1P

The structural properties of β-1,3-glucan significantly influence its biological functions and applications(Chen et al., 2022). To evaluate the β-1,3-glucan synthesized by TaβGP using G1P and glucose, its structure was characterized using SEM, XRD, FT-IR, and MALDI-TOF.

The β-1,3-glucan synthesized by TaβGP exhibited a pure white color (Fig. S5), in contrast to the off-white and faint yellow appearances of β-1,3-glucan products extracted from yeast, such as BYG and Glucan 300, respectively(Qiao et al., 2022). SEM analysis revealed that the synthesized β-1,3-glucan aggregated into homogeneous particles approximately 10 μm in diameter (Fig. 6A, 6B). Comparatively, BYG extracted using β-1,6-glucanase exhibited a relatively uniform size distribution of 1 × 5 μm(Qiao et al., 2022), while yeast β-glucan prepared via hot alkali-acetic acid treatment showed a heterogeneous size distribution(Hunter Jr et al., 2002). Both the synthesized β-1,3-glucan and Glucan 300 exhibited characteristic XRD diffraction peaks at 2*θ* = 6.5°and 20° (Fig. 6C). FT-IR analysis further confirmed their structural similarity, displaying analogous absorption spectra (Fig. 7D), including the characteristic β-configuration peak at 890 cm^−1^, and stretching vibration peaks corresponding to C=O, C-H, and OH groups at 1040, 2920, and 3420 cm^−1^, respectively(Liu et al., 2020). These results indicate that the synthesized β-1,3-glucan retains the structural integrity of natural β-1,3-glucan, which is essential for its bioactive properties.

**Fig. 6.**
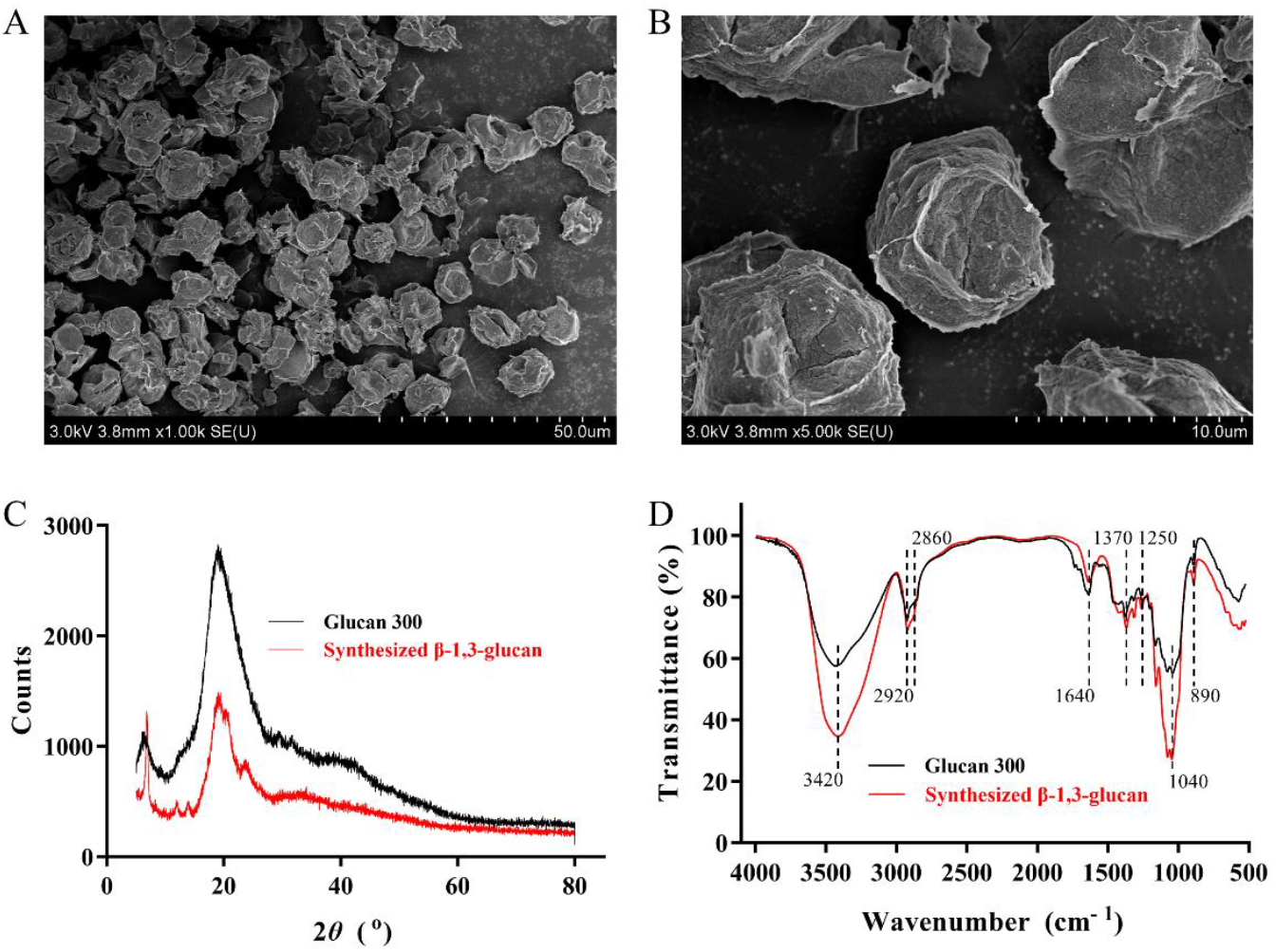
SEM (A, B), XRD (C), and FT-IR (D) of the β-1,3-glucan synthesized from glucose and G1P. The molecular wights of the synthesized β-1,3-glucan were determined by MALDI-TOF MS (Table S4). Previous studies(De Doncker et al., 2024; Zhong and Nidetzky, 2020) have reported that the DP of glucan can be modulated by the concentration of the priming substrate glucose. At 50 °C, TaβGP was capable of synthesizing β-1,3-glucan with a maximum DP of 43 (DP_n_ = 32.5) in the presence of 1 mM glucose. When 10 mM glucose were supplied, the resulting β-1,3-glucan had a DP_n_ of 25.7. Conversely, with 100 mM glucose, the synthesized β-1,3-glucan remained soluble, exhibiting a DP_n_ of 9.7.

## 4. Conclusions

In this study, THA_1941 from *T. africanus* (TaβGP) was re-identified as a GH61 β-1,3-glucan phosphorylase. TaβGP represents the first thermostable β-1,3-glucan phosphorylase in the GH161 family, retaining 100% of its original activity at 50 °C for 180 h. Notably, TaβGP’s unique ability to use glucose as a priming substrate eliminates the reliance on costly laminaribiose, addressing a major economic barrier in β-1,3-glucan production. The enzyme’s substrate flexibility, spanning α/β-linked disaccharides and diverse monosaccharides, coupled with resistance to metal ions (Na^+^, K^+^, Ca^2+^, Mg^2+^) and lignocellulose-derived inhibitors (e.g., furfural, 5-HMF), underscores its versatility for bioprocessing applications. Collectively, these findings establish TaβGP as a highly promising biocatalyst for industrial biomanufacturing, particularly for integration into *in vitro* enzymatic biosystems for the sustainable synthesis of β-1,3-glucan for use in pharmaceuticals, food additives, and biodegradable materials.

## CRediT authorship contribution statement

**Guotao Mao**: Conceptualization, Methodology, Writing – original draft, Writing – review & editing, Funding acquisition. **Jin Yu**: Methodology, Investigation, Writing – original draft. **Junhan Lin**: Investigation. **Ming Song**: Investigation. **Zengping Su**: Data curation, Supervision. **Hui Xie**: Data curation. **Hognsen Zhang**: Data curation. **Hongge Chen**: Conceptualization, Data curation, Writing – review & editing. **Andong Song**: Supervision, Writing – review & editing.

## Acknowledgments

This work was supported by the National Key Research and Development Program of China (No. 2022YFA3401700) and National Natural Science Foundation of China (No. 31800050).

## Appendix A. Supplementary data

The following is the Supplementary data to this article.

