## Supplementary Data for "THA_1941 from *Thermosipho africanus*: A Thermostable β-1,3-Glucan Phosphorylase for Efficient β-1,3-Glucan Synthesis"

### 1. Methods

Multiple sequence alignment of GH161 enzymes THA_1941 and PapP (NCBI accession No.: AUO30192.1), GH94 laminaribiose phosphorylase from *Paenibacillus* sp. (PsLBP, BAJ10826.1), and GH149 enzymes AtβOGP from *Anaerolinea thermo*phila (BAJ63785.1) and Pro_7066 (PDB: 6HQ8) was performed using Clustal Omega(Sievers and Higgins, 2018), and was further visualized using BioEdit(Hall et al., 2011).

### 2. Tables and figures

**Table S1. Primers for mutation construction**

| Primer | Sequence |
|---|---|
| R345Q-F | ATAACGTTTTACAAGGCGGTTATCCGCTGG |
| R345Q-R | GGATAACCGCCTTGTAAAACGTTATCCAGGTA |
| R373Q-F | CGGCGATCTGGAACAAGATTACAACTTCTTCG |
| R373Q-R | AAGTTGTAATCTTGTTCCAGATCGCCGTGTTT |
| D625A-F | CCGGGTTGGAACGCCGCATTAAACGGTTTA |
| D625A-R | CCGTTTAATGCGGCGTTCCAACCCGGTTTGTT |
| E840Q-F | CCCCGGGTTGGTTACAAAACGAGTCCGTCTTCAT |
| E840Q-R | CGGACTCGTTTTGTAACCAACCCGGGGTAAAT |

**Table S2. Chemical shifts (δ/ppm) of $^{13}C$ NMR signals for β-1,3-glucan**

| | C1 | C2 | C3 | C4 | C5 | C6 | Source |
|---|---|---|---|---|---|---|---|
| β-1,3-glucan | 103.0 | 72.8 | 86.2 | 68.4 | 76.3 | 60.8 | This work |
| β-1,3-glucan | 103.5 | 73.3 | 86.7 | 68.9 | 76.8 | 61.3 | (Zheng et al., 2016) |

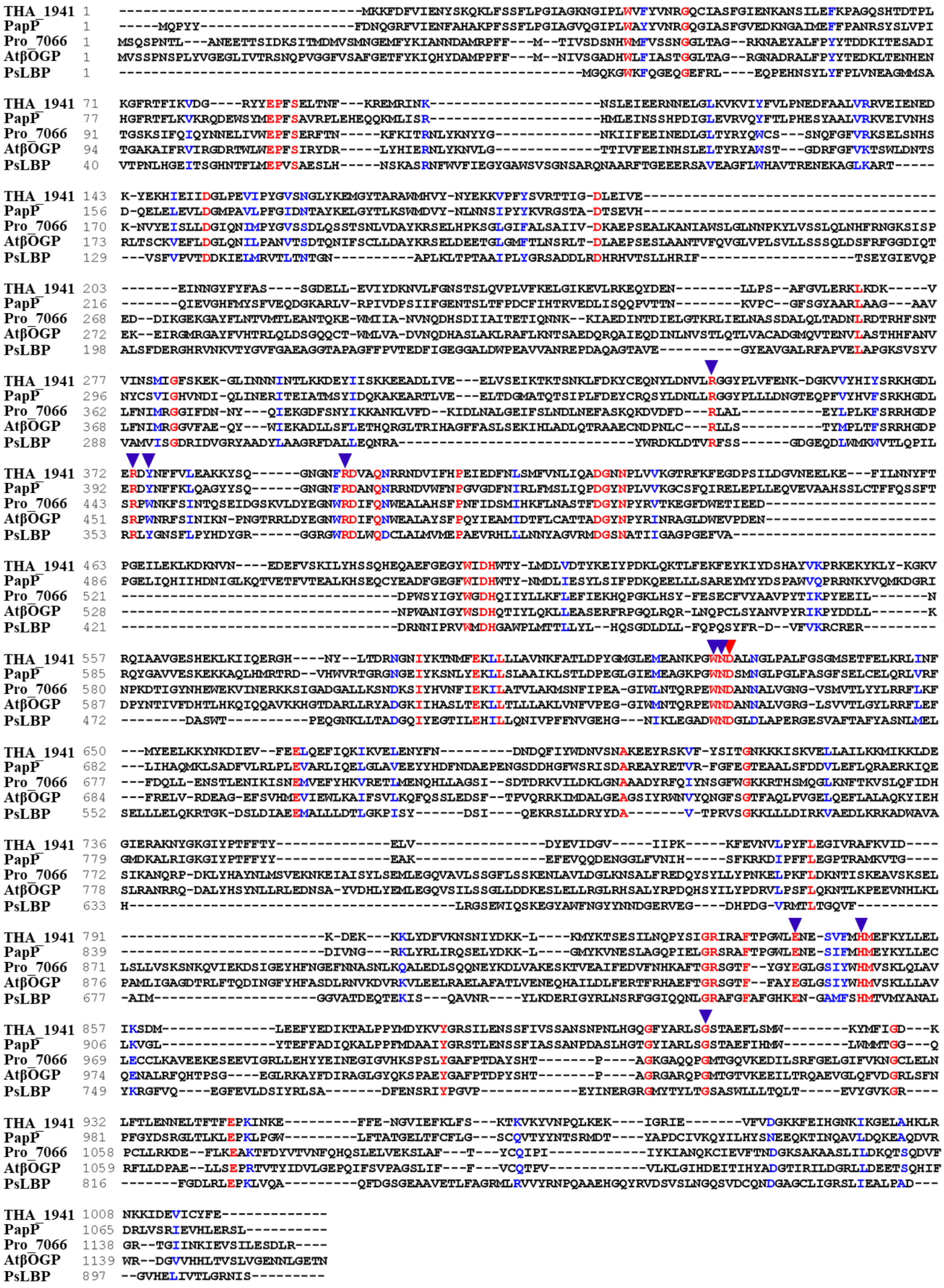

**Fig. S1. Sequence alignment of THA_1941 with other glycoside phosphorylases.**

The identical and similar residues were highlighted in red and blue, respectively. The conserved catalytic residue and glucose 1-phosphate binding residues were labelled with red and blue triangles.

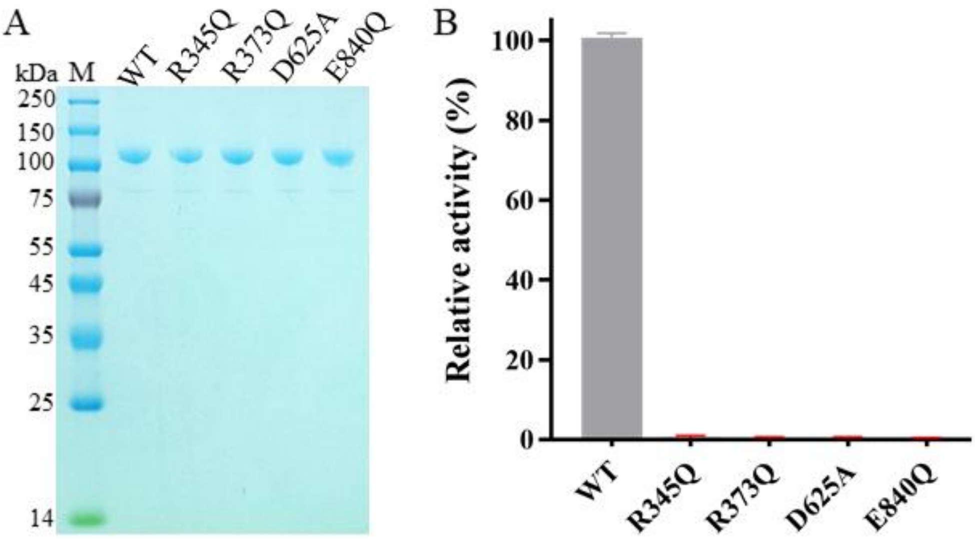


**Fig. S2. SDS-PAGE (A) and activity (B) of THA_1941 WT and mutants.** M and WT represented protein marker and THA_1941 wild type, respectively. The relatively activity of THA_1941 in the synthetic reaction was detected using glucose and glucose 1-phosphate.

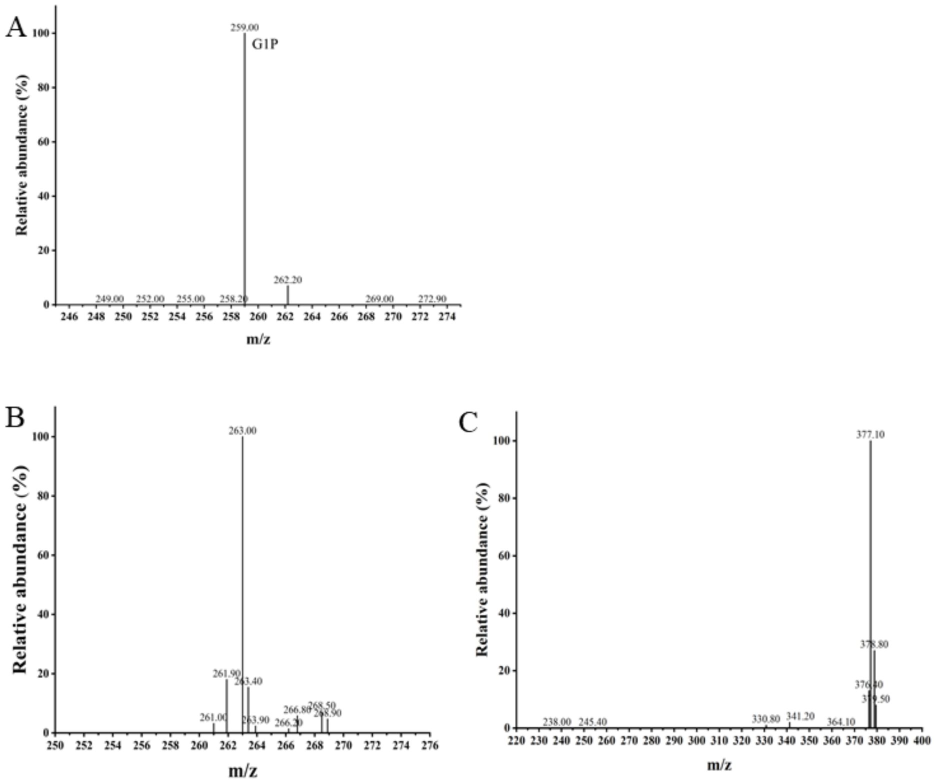


**Fig. S3. Analysis of G1P produced by THA_1941 in the phosphorolytic reaction by LC-MS.** (A) G1P standard. Detection of G1P produced in the phosphorolytic reaction utilizing cellobisoe (B) and cellotriose (C) as the substrate.

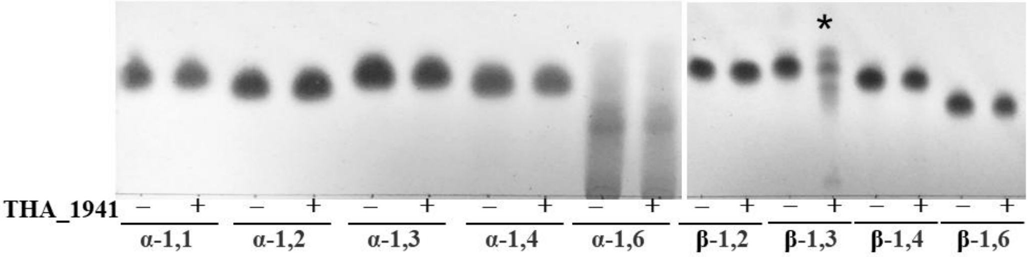


**Fig. S4. TLC analysis of the phosphorolytic reaction catalyzed by THA_1941 utilizing glucose-glucose disaccharides.**

The linkages of the disaccharides were indicated. The effective substrates of THA_1941 detected by TLC were marked with asterisks.

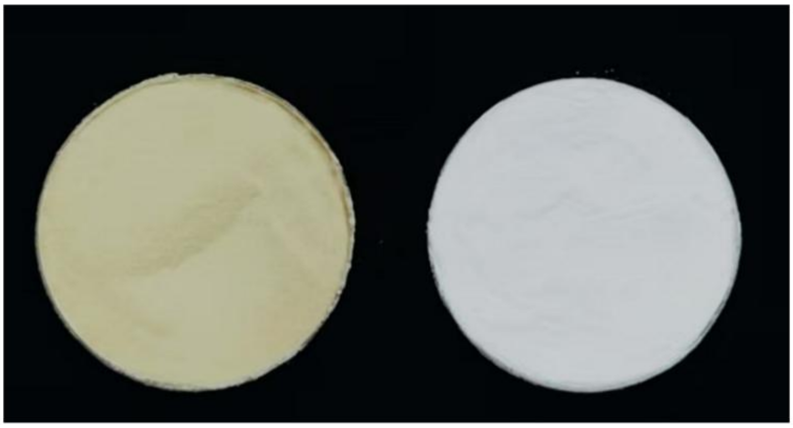


**Fig. S5. Macroscopical appearance of Glucan 300 and the synthesized β-1,3-glucan using glucose and G1P**
